# Facilitation of motor excitability during listening to spoken sentences is not modulated by noise or semantic coherence

**DOI:** 10.1101/168864

**Authors:** Muriel T.N. Panouillères, Rowan Boyles, Jennifer Chesters, Kate E. Watkins, Riikka Möttönen

## Abstract

Comprehending speech can be particularly challenging in a noisy environment and in the absence of semantic context. It has been proposed that the articulatory motor system would be recruited especially in difficult listening conditions. However, it remains unknown how signal-to-noise ratio (SNR) and semantic context affect the recruitment of the articulatory motor system when listening to continuous speech. The aim of the present study was to address the hypothesis that involvement of the articulatory motor cortex increases when the intelligibility and clarity of the spoken sentences decreases, because of noise and the lack of semantic context. We applied Transcranial Magnetic Stimulation (TMS) to the lip and hand representations in the primary motor cortex and measured motor evoked potentials from the lip and hand muscles, respectively, to evaluate motor excitability when young adults listened to sentences. In Experiment 1, we found that the excitability of the lip motor cortex was facilitated during listening to both semantically anomalous and coherent sentences in noise, but neither SNR nor semantic context modulated the facilitation. In Experiment 2, we replicated these findings and found no difference in the excitability of the lip motor cortex between sentences in noise and clear sentences without noise. Thus, our results show that the articulatory motor cortex is involved in speech processing even in optimal and ecologically valid listening conditions and that its involvement is not modulated by the intelligibility and clarity of speech.

## 1. Introduction

Speech perception is a demanding skill that is supported by an extensive brain network. Although the human auditory system is critical for the processing of acoustic speech signals, numerous neuroimaging studies have shown that frontal cortical regions such as inferior frontal gyrus (IFG) and premotor cortex are also activated during speech perception (Adank, 2012; Callan, Callan, Gamez, Sato, & Kawato, 2010; Hervais-Adelman, Carlyon, Johnsrude, & Davis, 2012; Londei et al., 2010; Osnes, Hugdahl, & Specht, 2011; Pulvermüller et al., 2006; Skipper, Devlin, & Lametti, 2017; Skipper, Nusbaum, & Small, 2005; Szenkovits, Peelle, Norris, & Davis, 2012; Wilson, Saygin, Sereno, & Iacoboni, 2004). Transcranial magnetic stimulation (TMS) combined with electromyography provides a method to measure excitability of the representations of the articulators in the primary motor cortex during speech perception (Adank, Nuttall, & Kennedy-Higgins, 2017; Möttönen, Rogers, & Watkins, 2014; Möttönen & Watkins, 2012). Single TMS pulses over the representations of the articulators in the primary motor cortex elicit motor evoked potentials (MEPs) in the targeted muscles. Changes in the size of MEPs reflect changes in the excitability of the motor pathways connecting the cortical representations with the corresponding muscles. Using this technique, several studies have demonstrated that the excitability of the primary motor cortex, controlling articulatory gestures to produce speech, is enhanced during listening to speech (Fadiga, Craighero, Buccino, & Rizzolatti, 2002; Murakami, Restle, & Ziemann, 2011; Murakami, Ugawa, & Ziemann, 2013; Nuttall, Kennedy-Higgins, Devlin, & Adank, 2017; Nuttall, Kennedy-Higgins, Hogan, Devlin, & Adank, 2016; Watkins, Strafella, & Paus, 2003).

It has been proposed that the articulatory motor system is a complementary system, recruited when listening to speech in challenging conditions (Wilson, 2009). Some MEP studies have indeed shown that listening to speech in noise enhances the excitability of the lip motor cortex more than listening to speech (sentences or syllables) without noise (Murakami et al., 2011; Nuttall et al., 2017). These MEP studies did not however include a wide range of noise levels and therefore it is currently unknown how signal-to-noise ratio (SNR) of speech signal affects the excitability of the articulatory motor cortex. Several functional Magnetic Resonance Imaging (fMRI) studies have found an increased activation in the speech motor system to degraded speech compared to clear speech (Adank & Devlin, 2010; Du, Buchsbaum, Grady, & Alain, 2014; Evans & Davis, 2015; Hervais-Adelman et al., 2012; Osnes et al., 2011). Recently, Du et al (2014) investigated the activation of the motor and auditory systems during a phoneme categorization task at various SNR levels. The activation of the speech motor system (premotor cortex and posterior IFG) correlated negatively with the SNR-modulated accuracy. Furthermore, multi-voxel pattern analyses showed that the speech motor cortex successfully categorized the phonemes at lower SNR levels than the auditory system. These findings support the idea that the speech motor system has a compensatory role when categorizing speech sounds in noisy conditions. However, since the participants performed a button press task, it is unclear whether the motor activations were related to this task or processing of speech sounds (see for a discussion of this point Schomers & Pulvermüller, 2016). Importantly, it remains unknown how SNR affects the activity of the articulatory motor system during passive listening to more natural speech signals such as sentences.

In everyday life, speech comprehension is supported by semantic context as it improves intelligibility of continuous speech in noise (Davis, Johnsrude, Hervais-Adelman, Taylor, & McGettigan, 2005; Miller & Isard, 1963; Obleser, Wise, Dresner, & Scott, 2007). For example, word report scores for semantically coherent sentences like “the coin was thrown onto the floor” are higher than for semantically anomalous sentences like “the boot was grown onto the mouth” across a wide range of SNR levels (Davis, Ford, Kherif, & Johnsrude, 2011). Neuroimaging studies have shown that semantic context affects activity in the IFG and its connectivity with other brain regions (Davis et al., 2011, 2005; Obleser et al., 2007; Sohoglu, Peelle, Carlyon, & Davis, 2012). It can be hypothesized that if the articulatory motor system contributes to speech perception in challenging conditions, then it should show increased activation when listening to semantically anomalous sentences relative to semantically coherent sentences. Whether the excitability of the articulatory motor system is modulated by semantic coherence of spoken sentences remains to be investigated.

In the present study, we aimed to address the hypothesis that the recruitment of the articulatory motor cortex increases when the intelligibility of the spoken sentences decreases and speech perception becomes more challenging. We modulated intelligibility of spoken sentences by manipulating their SNR and semantic coherence. MEPs from the lip and the hand muscles were measured while participants passively listened to semantically coherent and anomalous sentences and non-speech signals in two experiments. The aim of Experiment 1 was to test how a range of five SNR levels affects motor excitability. The aim of Experiment 2 was to test replicability of the results of Experiment 1 and to determine whether motor excitability is sensitive to the presence of noise when processing spoken sentences. Experiment 2 included sentences at two SNR levels and sentences without noise. The comparison between lip and hand MEPs allowed us to test whether listening to speech enhances excitability in the articulatory motor system specifically.

## 2. Materials and Methods

### 2.1. Participants

Forty participants were recruited in Experiment 1. The data of 11 participants were excluded because of 1) unreliable motor evoked potentials (MEPs) in the lip muscle (N=4), 2) artefacts in the recording preventing the accurate offline detection of lip MEPs (N=5), 3) lip background muscle contraction (N=1) and 4) proportion of correctly reported words for the anomalous sentences was below 40% at the highest SNR (0dB) (N=1). In total, we report the data from 29 participants for Experiment 1. Thirteen participants were in the hand group (7 females –age: 24.4 ±5.3 years old) and sixteen in the lip group (5 females – age: 22.9 ±3.9 years old).

Thirty-five participants were recruited in Experiment 2. The data of 10 participants was excluded based on 1) lip background muscle contraction (N=8), 2) artefacts in the recording preventing the accurate offline detection of lip MEPs (N=1) and 3) the reported clarity in the anomalous sentences at the highest SNR (0dB) was below 3.2 (equivalent to 40% reported accuracy in Experiment 1; N=1). In total, we report the data from 16 participants in the hand group (10 females –age: 22.4 ±2.8 years old) and 9 participants in the lip group (six females –age: 24.4 ±3.6 years old).

The 54 participants for whom data is reported in the present study are right-handed, native-English speakers, with no known neurological, psychiatric, hearing or language impairment. All participants gave their written informed consent and were screened prior inclusion for contraindications to TMS. Experimental procedures conformed to the Code of Ethics of the World Medical Association (Declaration of Helsinki) and were approved by Oxfordshire NHS Research Ethics Committee B (REC Reference Number 10/H0605/7).

### 2.2. Electromyography

Electromyography (EMG) activity was recorded using surface electrodes (22 × 30 mm ARBO neonatal electrocardiogram electrodes). Recordings from the right orbicularis oris were taken from electrodes attached to the right upper and lower lip. Recordings from the right first dorsal interosseous muscle were taken from electrodes attached to the belly and tendon of the muscle. The ground electrode was attached to the forehead. The raw EMG signal was amplified (gain: 1000), bandpass filtered (1-1000Hz) and sampled (5000Hz) via a CED 1902 four-channel amplifier, a CED 1401 analog-to-digital converter and a computer running Spike2 (Cambridge Electronic Design). The EMG signals were stored on the computer for off-line analysis.

### 2.3. Transcranial Magnetic Stimulation

All TMS pulses were monophasic, generated by Magstim 200 (Magstim, Whitland, UK) and delivered through a 70-mm figure of eight coil. The position of the coil over the left motor cortex was adjusted until a robust motor-evoked potential (MEP) was observed in the contralateral target muscle (either hand or lip). Single-pulse TMS was delivered for every trial to allow recording MEPs from the resting target muscle. The intensity of the stimulation was set in order for an MEP of at least 1mV peak-to-peak for the hand and at least 0.2mV for the lip to be consistently produced in the resting muscle. The mean intensity used for the lip groups was of 69.8% (±6.9%) and of 58.2% (±8.2%) in Experiments 1 and 2, respectively. For the hand stimulation, the averaged intensity was of 59.7% (±12.0%) and of 56.5 (±9.5%) in Experiments 1 and 2, respectively.

### 2.4. Stimuli

The stimuli used in the present study have been used in previous fMRI studies (Davis et al., 2011; Rodd, Davis, & Johnsrude, 2005). The set comprised 200 declarative sentences between 6 and 13 words in length. One hundred sentences from this set were semantically coherent. The remaining hundred sentences were semantically anomalous created by randomly substituting content words matched for syntactic class, frequency of occurrence and number of syllables. The anomalous sentences were identical to the normal sentences in terms of phonological, lexical and syntactic properties but lacked coherent meaning. All 200 sentences (1.2 to 3.5 seconds in duration, speech rate 238 words/minute) were produced by a male speaker of British English and digitized at a sampling rate of 44.1Khz. The sentences were degraded by adding speech-spectrum signal-correlated noise (SCN) at a range of signal to noise ratios (SNRs) using Praat software (Davis et al., 2011). At all SNRs, duration, amplitude and average spectral composition of the sentence remained the same. The SNRs used in the present study were clear speech (no added noise), 0dB, -1dB, -2dB, - 3dB and -4dB. Pure SCN stimuli and white noise (WN) stimuli were also used in the present study. The pure SCN stimuli have a similar rhythmic structure as speech and therefore provide an acoustically matched non-speech baseline.

### 2.5. Experimental set-up and procedures

In both experiments, participants were either assigned to the lip or the hand stimulation group. Participants sat in front of a computer presenting the stimuli using Presentation^®^ software (Neurobehavioral Systems, Inc., Berkeley, CA, USA). Audio stimuli were presented to the participants through insert earphones (Etymotic, Elk Grove Village, IL, USA).

#### 2.5.1. Experiment 1

This experiment included two blocks, one with coherent sentences and the other one with anomalous sentences. In each block, 100 different sentences were presented, 20 of each at the SNR of 0dB, -1dB, -2dB, -3dB and -4dB, along with 30 SCN stimuli and 30 WN stimuli. For each block, the order of the 160 stimuli was randomized and the order of the blocks (coherent and anomalous) was counterbalanced across participants. Participants were instructed to listen to the sentences or to the noise while keeping both their lip and hand muscles relaxed. For each stimulus (sentence, SCN or WN), a single-pulse of TMS was delivered to elicit an MEP. For the sentence stimuli, it was delivered 150 ms after the onset of the final content word. This was chosen as a reliable way of matching the point at which TMS was delivered across sentences, as the final content word was likely to be the most predictable. For the WN and SCN stimuli, the pulse was delivered close to the end of the stimuli, matching the timing of the pulses for sentence stimuli. The average inter-pulse-interval was 6s (range: 4.24s–7.94s). Within each block, there was a short break every 32 trials.

After the completion of the two blocks, participants listened to the 100 anomalous and 100 coherent sentences again and repeated them out loud. For each type of sentences, 20 sentences were presented at the 5 SNR (0dB, -1dB, -2dB, -3dB and -4dB). The experimenter assessed the accuracy of the participant’s response during the task.

#### 2.5.2. Experiment 2

This experiment included only one session. Subjects were presented with 40 clear speech stimuli, 40 at 0dB and 40 at -2dB from the set of sentences described earlier. Half of the sentences were anomalous sentences and half of them were coherent sentences. The experiment included also 30 WN and 30 SCN stimuli. All stimuli were presented in random order. As in Experiment 1, a single TMS pulse was delivered in the beginning of the last word of the sentence (as above) eliciting an MEP in either the relaxed hand or the relaxed lip muscle.

After the TMS session, participants listened to a subset of the sentences again (90 in total, 15 of each SNR for the normal sentences, 15 of each SNR for the anomalous sentences) and were asked, after hearing each sentence, to rate the clarity of the speech on a scale from 1 to 8, using the computer keyboard.

### 2.6. Behavioural analysis

In Experiment 1, we calculated the proportions of correctly reported words for each SNR level and sentence type in each participant. In Experiment 2, we calculated means of clarity scores for each SNR level and sentence type in each participant.

### 2.7. MEP analysis

MEPs were analysed on a trial-by-trial basis using in-house software written in Matlab (Mathworks Inc, Natick, USA). Maximal and minimal peaks of the MEPs were automatically detected using a fixed window following the TMS pulse: [15-40ms] for the hand and [12-35ms] for the lip. The detection was checked manually by the experimenter. The absolute value of the background muscle activity was averaged across the 100ms preceding the TMS pulse and trials with a mean absolute value of background muscle activity higher than 2 standard deviations of the average for each TMS session were excluded. Outliers MEPs with values above or below 2 standard deviations of the mean for each experimental condition (sentence types and SNR levels) were removed. After removing outliers, we calculated MEP z-scores for each experimental condition relative to the WN in each participant.

### 2.8. Statistical analysis

Statistical analyses were performed with the SPSS Statistics software package (IBM, Armonk NY, USA). The proportion of correctly reported words, the perceived clarity of the sentences and the normalised MEP z-scores were analysed using separate ANOVAs with the within-subject factors semantic coherence (coherent vs anomalous) and SNR (Experiment 1: 0 dB, -1dB, -2 dB, -3 dB, -4 dB; Experiment 2: clear; 0dB, -2dB) and the between-subjects factor group (hand vs lip). In both experiments, because of a lack of SNR effect and semantic effect and interaction involving these factors, the MEP z-scores were averaged across SNR levels and semantic types. In Experiments 1 and 2, these averaged z-scores were submitted to an ANOVA with the within-subject factor stimulus (speech vs SCN) and the between-subjects factor group (hand vs lip).

For all ANOVAs, Greenhouse–Geisser corrections to the degrees of freedom were applied if Mauchly’s sphericity test revealed a violation of the assumption of sphericity for any of the factors in the ANOVAs. Significance level was set at P < 0.05.

## 3. Results

### 3.1. Experiment 1

The aim of Experiment 1 was to examine whether and to what extent SNR and semantic coherence of sentences modulate motor excitability. We measured MEPs from the lip and hand muscles while participants passively listened to sentences. After MEP measurements, we tested how SNR and semantic coherence affected the intelligibility of the sentences using a behavioural word-report task.

#### 3.1.1. Effect of semantic coherence and SNR on intelligibility of spoken sentences

Figure 1 presents the proportions of correctly reported words for the semantically coherent and anomalous sentences at the various SNRs (0 dB, -1dB, -2 dB, -3 dB, -4 dB). The proportion of correctly reported words decreased with the SNR (main effect of SNR: F[2,67]=581.73, p<0.001). Moreover, the intelligibility of the anomalous sentences was lower than intelligibility of the coherent sentences (main effect of semantic coherence: F[1,27]=141.31, p<0.001). An interaction between the semantic coherence and SNR was also significant (F[2,67]=18.01, p<0.01), mostly because the difference between anomalous and coherent sentences was greater at the intermediate SNRs (-1 to -3dB) than at other SNRs. No significant main effect of TMS group and interactions involving this factor were found (F[2,67]<1, p>0.55). Thus, SNR levels and semantic coherence modulated intelligibility similarly in the hand and lip groups.

**Figure 1:**
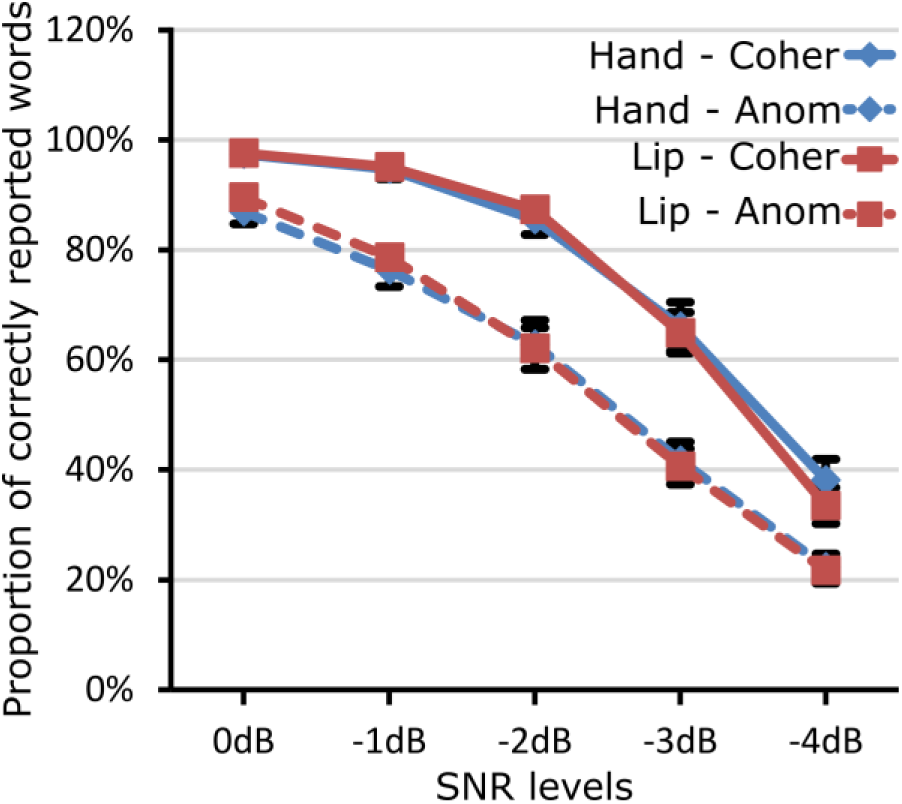
Behavioural performances in Experiment 1. Effects of SNR levels and sentence coherence on word-report accuracy in Experiment 1. The proportion of correctly reported words is represented as a function of the SNR levels for the hand (blue diamonds) and lip (red squares) groups, separately for the coherent (Coher: continuous line) and anomalous (Anom: dashed lines) sentences. Error bars are standard error of the mean.

#### 3.1.2. Motor excitability when listening to sentences

The MEP z-scores normalised to the WN baseline are presented for anomalous and coherent sentences at the five SNR levels in Figures 2A and 2B for the lip and hand groups, respectively. To test whether SNR and semantic affected motor excitability, a three-way ANOVA for the MEP z-scores with SNR and semantic coherence as within-subject factors and group as a between-subjects factor was carried out. There was no significant main effect or interaction involving the SNR factor or the semantic coherence factor (F[4,108]<1.30, p>0.27), suggesting that motor excitability was stable across the five SNR levels and across the sentence types. The z-scores for all speech stimuli (across five SNR levels and coherent and anomalous sentences) were then averaged for each participant in order to examine whether listening to speech enhanced motor excitability relative to non-speech stimuli.

**Figure 2:**
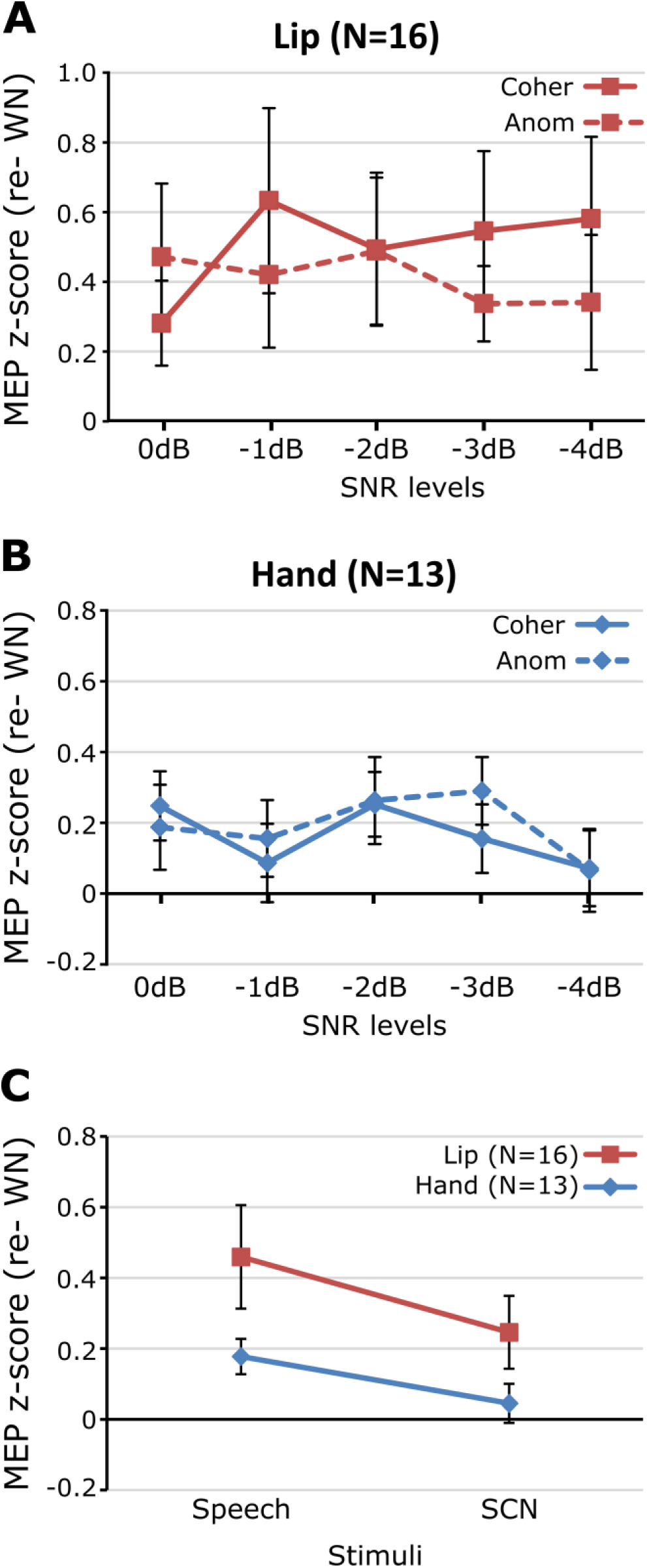
MEPs of the lip and hand groups in Experiment 1. MEP z-scores during the perception of sentences in noise and speech-correlated noise (SCN). The MEPs elicited during the perception of sentences in the five SNR levels are represented for the lip (A) and hand group (B), separately for the coherent (Coher) and Anomalous (Anom) sentences. These z-scores are shown averaged for the speech stimuli and compared to the non-speech stimuli (SCN) for the lip (red squares) and hand (blue diamonds) groups (C). Error bars are standard error of the mean.

Figure 2C presents the MEP z-scores for the speech and SCN stimuli in the lip and hand groups. To assess whether the stimulus type modulated motor excitability, an ANOVA with the within-subject factor stimulus type (speech vs SCN) and the between-subjects factor TMS group (hand vs lip) was performed. The main effect of the stimulus type was significant (F[1,27]=16.96, p<0.001), showing that motor excitability was greater when listening to speech than when listening to SCN. There was no significant main effect of the group factor nor any interaction between group and stimuli (F[1,27]<2.08, p>0.16).

To assess whether motor excitability was enhanced relative to the WN baseline, the MEP z-scores (normalized relative to WN), were compared statistically to 0. The lip MEP z-scores were significantly greater than 0 for the speech stimuli (one sample t-tests: t[15]=2.67, p<0.05) and was slightly enhanced for the SCN (one sample t-tests: t[15]=2.03, p=0.06). The hand MEP z-scores were greater than 0 only for the speech stimuli (t[12]=3.42, p<0.01) but not for the SCN (t[12]=0.79, p=0.45).

In summary, in Experiment 1 we found no modulatory effect of SNR or semantic coherence on motor excitability during listening to spoken sentences. As expected, listening to speech however enhanced the excitability of the lip motor cortex relative to non-speech sounds (WN and SCN). Unexpectedly, the excitability of the hand motor cortex was also enhanced during listening to speech relative to non-speech sounds.

### 3.2. Experiment 2

In Experiment 1, we found no effect of SNR on the motor excitability when participants listened to spoken sentences. However, all sentences were presented in noise. In Experiment 2, we examined whether the presence of noise can enhance motor excitability by using sentences with (SNRs: 0 and - 2dB) and without noise (clear speech). Furthermore, in Experiment 1 we presented anomalous and coherent sentences in different blocks while MEPs were recorded from the lip and hand muscles, which may have reduced the reliability of the comparison between sentences types. In Experiment 2, we presented anomalous and coherent sentences in the same block in order to examine the effect of semantic coherence on motor excitability more reliably. Similarly to Experiment 1, two non-speech stimuli were included in the block (WN baseline and SCN) and participants were either assigned to the hand or to the lip group. After MEP measurements, we tested how the presence of noise and semantic coherence affect the perceived clarity of the spoken sentences using a rating task.

#### 3.2.1. Effect of semantic coherence and SNR on perceived clarity of spoken sentences

Figure 3 presents the mean clarity ratings for the anomalous and coherent sentences presented without noise and with SNRs of 0dB and -2dB. Similarly to Experiment 1, the main effects of semantic coherence (F[1,23]=107.23, p<0.001) and SNR (F[1, 34]=157.66, p<0.001) as well as an interaction between semantic coherence and SNR (F[2,46]=44.43, p<0.001) were significant. This demonstrates that the anomalous sentences were less clear than the coherent sentences at the 0dB and -2dB SNR levels, whereas there was no difference in clarity in the absence of noise between the two sentences types. There was a non-significant tendency for lower clarity ratings in the lip group than in hand group (F[1,23]<3.35, p>0.08), but no significant interactions involving TMS group. In sum, SNR and semantic coherence modulated the perceptual clarity of the sentences in both groups.

**Figure 3:**
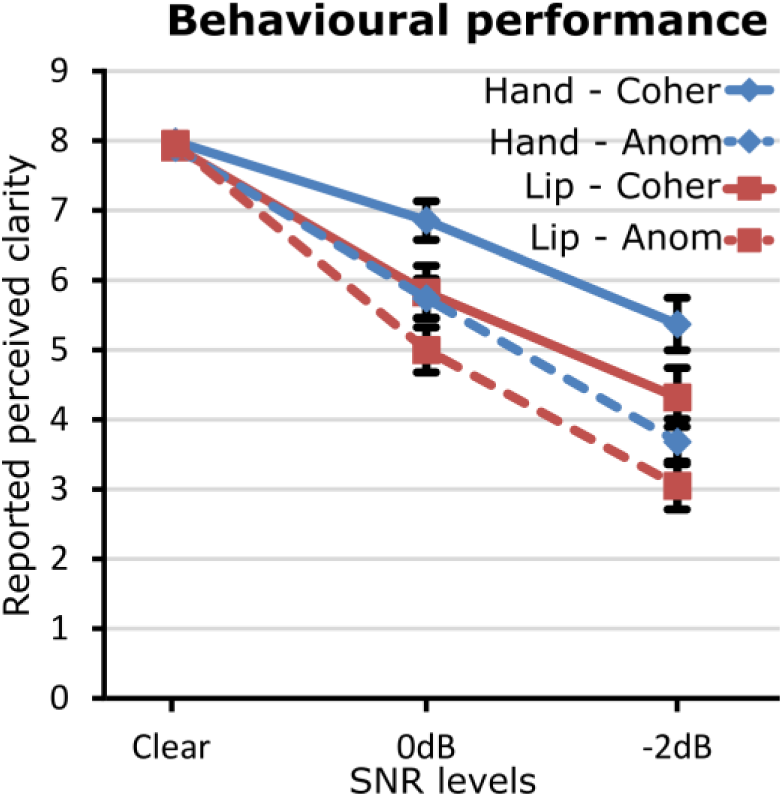
behavioural performance in Experiment 2. Effects of the presence of noise and sentence coherence on reported perceived clarity in Experiment 2. The perceived clarity is represented as a function of the SNR levels for the hand (blue diamonds) and lip (red squares) groups, separately for the coherent (continuous line) and anomalous (dashed lines) sentences. Error bars are standard error of the mean.

#### 3.2.2. Specific facilitation of the lip motor cortex when listening to speech

The MEP z-scores, normalised to the WN baseline, for anomalous and coherent sentences are presented as a function of SNR in Figures 4A and 4B for the lip and hand groups, respectively. To evaluate whether the presence of noise and semantic coherence modulates motor excitability during listening to speech, an ANOVA with the within-subject factors SNR (clear vs 0dB vs -2dB) and semantic coherence (coherent vs anomalous) and the between-subjects factor TMS group (hand vs lip) was carried out. There was no significant main effect of SNR, semantic coherence or interactions involving these factors (F[2,46]<2.22, p>0.12), suggesting that motor excitability was not modulated by the presence of noise nor by semantic coherence. The main effect of the TMS group was significant (F[1,23]=3.24, p<0.05) showing that listening to speech enhanced the excitability of the lip motor cortex more than the excitability of the hand motor cortex. Because the SNR and semantic coherence factors had no effect, we averaged the MEP z-scores across the three SNR levels and the two sentence types for each participant (Figure 4C). To test whether motor excitability was modulated by stimulus type, an ANOVA with the within-subjects factor stimulus (speech vs SCN) and the between-subjects factor group (hand vs lip) was carried out on the MEP z-scores. Stimulus type differently modulated the excitability of the lip and hand motor cortex (stimulus effect: F[1,23]=3.99, p=0.06; stimulus × group interaction: F[1,23]=10.47, p<0.01). Pairwise comparisons revealed that listening to speech stimuli enhanced the excitability of the lip motor cortex relative to SCN (t[8]=4.1, p<0.01) but that the excitability of the hand motor cortex was not modulated by the stimulus type (i.e., speech vs. non-speech - t[15]=-0.95, p=0.36).

**Figure 4:**
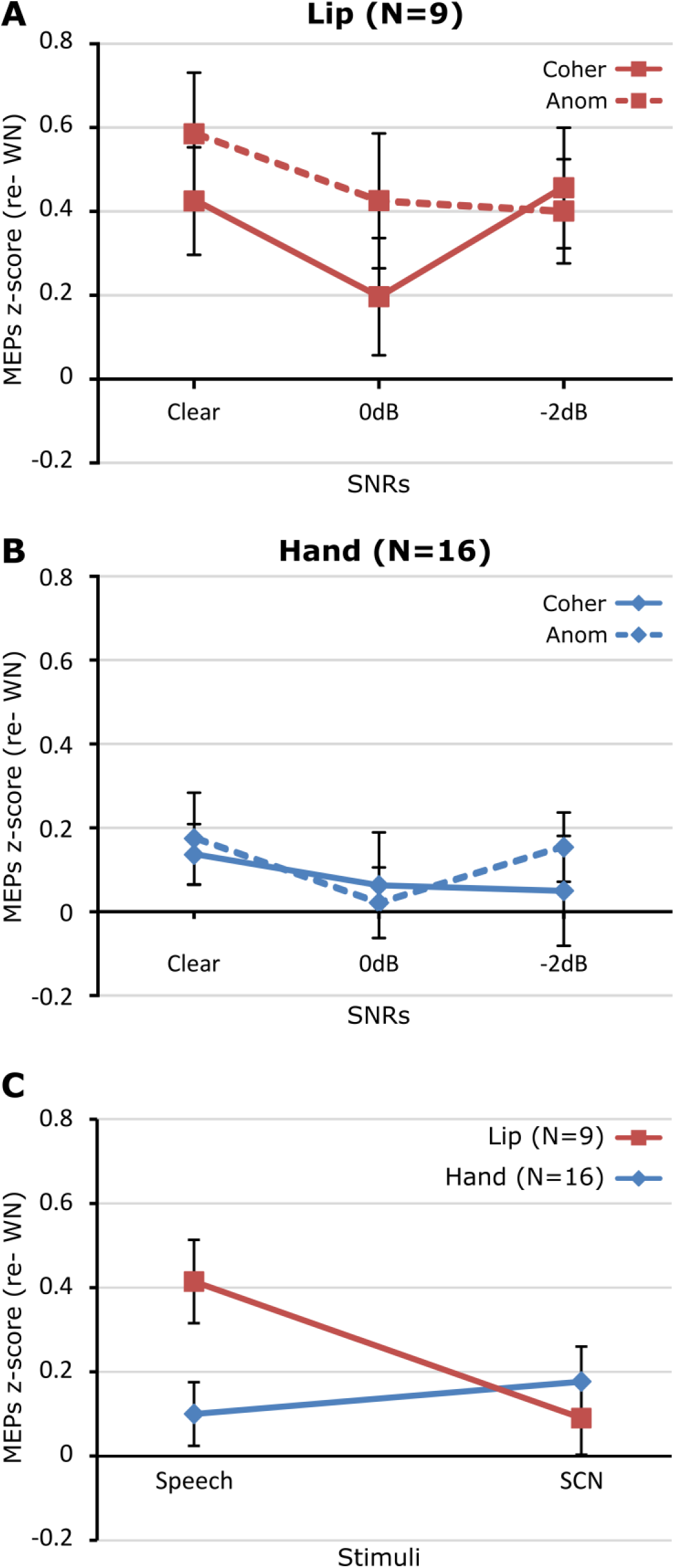
MEPs of the lip and hand groups in Experiment 2. MEP z-scores during the perception of sentences with and without noise and speech-correlated noise (SCN). The MEPs elicited during the perception of sentences in clear speech, at 0dB and -2dB are represented for the lip (A) and hand group (B), separately for the coherent (Coher) and Anomalous (Anom) sentences. These z-scores are represented averaged across the speech stimuli and for the non-speech stimuli (SCN) for the lip (red squares) and hand (blue diamonds) groups (C). Error bars are standard error of the mean.

To assess whether motor excitability was enhanced relative to the WN baseline, the MEP z-scores (normalized relative to WN) were statistically compared to 0. The lip MEP z-scores were significantly greater than 0 for the speech stimuli (one sample t-test: t[8]=4.19, p<0.01) but not for the SCN (t[8]=1.04, p=0.33). In the hand group, the MEP z-scores for the speech stimuli did not differ from 0 (t[15]=1.32, p=0.21), but they were marginally greater than 0 for the SCN (t[15]=2.12, p=0.05).

In summary, results of Experiment 2 showed that passive listening to spoken sentences enhances the excitability of the lip motor cortex more than listening to non-speech sounds (SCN and WN), and that neither semantic coherence of the sentences nor SNR (0 vs -2 dB) affect the excitability of the lip motor cortex. These findings replicate the findings of Experiment 1. Furthermore, we found no evidence that the presence of noise modulates the excitability of the lip motor cortex, since no differences were found between clear sentences and sentences presented in noise. Moreover, excitability of the hand motor cortex was not modulated by listening to speech stimuli (relative to SCN and WN).

## 4. Discussion

In this study, we aimed to address the hypothesis that the involvement of the articulatory motor cortex in speech processing increases when speech is difficult to understand. We manipulated the intelligibility and clarity of spoken sentences by modulating their SNR and semantic coherence. Results of Experiments 1 and 2 showed that listening to spoken sentences increased the excitability of the lip motor cortex more than listening to non-speech signals. Changes in excitability in the hand motor cortex were smaller and less consistent across experiments. Importantly, SNR and semantic coherence had no influence on the excitability of the lip motor cortex in either experiment. Thus, we found no supporting evidence for the hypothesis that the involvement of the articulatory motor cortex increases in challenging listening conditions. Our findings show that the articulatory motor cortex is involved in speech processing even in optimal and ecologically valid listening conditions and that its involvement is not modulated by the intelligibility and clarity of speech.

In both Experiments 1 and 2, listening to speech enhanced the excitability of the lip motor cortex relative to non-speech signals (i.e., WN and SCN). This shows that the articulatory motor cortex is involved in speech processing in agreement with previous studies (Fadiga et al., 2002; Murakami et al., 2011; Watkins et al., 2003). Our experiments included two non-speech signals, WN and SCN, which have different temporal characteristics. WN is stationary, whereas SCN has speech-like temporal structure. In Experiment 1, listening to SCN slightly enhanced lip excitability relative to WN, while no such enhancement was found in Experiment 2. The main difference between the two experiments was that Experiment 1 included lower SNRs level (-3dB and -4dB) than Experiment 2 (clear speech to -2 dB). It is possible that the participants were uncertain about the presence of speech signal in SCN in the Experiment 1, but not in Experiment 2, explaining the enhanced excitability of the lip motor cortex in Experiment 1 for SCN. Thus, although listening to speech signals enhances the excitability of the lip motor cortex more than listening to non-speech signals, the excitability can be slightly enhanced during listening to dynamic non-speech signals in some circumstances. However, this enhancement was not specific to the articulatory motor cortex as the excitability of the hand motor cortex was also enhanced when listening to SCN relative to WN in Experiment 2. Moreover, the excitability of the hand motor cortex was enhanced during listening to speech relative to WN in Experiment 1, but this was not replicated in Experiment 2. Thus, no consistent speech-specific enhancements in the excitability of the hand motor cortex were found in Experiments 1 and 2.

Our behavioural results showed that the semantically coherent sentences were more intelligible (Experiment 1) and clearer (Experiment 2) than semantically anomalous sentences, replicating findings from previous studies (Davis et al., 2005; Miller & Isard, 1963). We hypothesized that if the articulatory motor system is involved in speech perception especially in challenging conditions (Wilson, 2009), the excitability should be enhanced more during listening to semantically anomalous sentences than semantically coherent sentences. We found no support for this hypothesis as listening to coherent and anomalous sentences equally facilitated the excitability of the lip motor cortex.

We also manipulated the difficulty of speech perception by varying the SNR of the sentences. The SNR had a strong effect on intelligibility (Experiment 1) and perceived clarity (Experiment 2) of the spoken sentences. Despite this, no difference in excitability of the lip motor cortex was found between the five levels of SNR in Experiment 1 and between clear speech and speech in noise in Experiment 2. In contrast, two earlier studies have demonstrated an increase of lip motor excitability when listening to speech in noise relative to clear speech (Murakami et al., 2011; Nuttall et al., 2017). Murakami et al (2011) measured the lip motor excitability when participants listened to clear sentences or sentences in white noise (their Experiment 4). In terms of behaviour, participants were able to accurately report the clear sentences (^~^100% accuracy) while the addition of white noise decreased intelligibility by approximately 20% (^~^80% accuracy). Listening to both types of sentences facilitated lip motor excitability relative to the white noise baseline, but the facilitation was greater for sentences in noise than clear sentences. Nuttall et al (2017) tested the effect of speech-shaped unmodulated noise on lip motor excitability during speech perception in two experiments. In the first experiment, participants listened to four different vowel-consonant-vowel (VCV) syllables with or without noise. The amount of noise added to the syllables varied across participants in order to match another experimental condition. The accuracy report of the syllables in noise varied from 15% to 87.5% across individuals and this manipulation of intelligibility enhanced the lip motor excitability relative to the clear speech condition. In the second experiment, participants listened to VCVs at two different levels of noise with accuracy of 50% and 75%. No differences in lip motor excitability were found between speech in noise and clear speech in this second experiment. The authors explained this apparent discrepancy between their two experiments by the fact that the listening conditions were on average more challenging in the first than in the second experiment. With our SNR manipulation, we modulated speech intelligibility from 20% to 100% accuracy, but did not find a further facilitation of the lip motor cortex for the more challenging listening conditions relative to the easy listening conditions. Both Murakami et al (2011) and Nuttall et al (2017) repeated the same stimuli several times during their TMS experiments, whereas sentences were never repeated during the MEP recordings in the present study. This stimulus repetition may then have an effect on lip motor excitability. Thus, differences in the type of noise (white noise versus speech-correlated noise), in the type of speech stimuli (sentences versus syllables) and in stimulus repetition could potentially explain the differences with the present results and previous ones. Nevertheless, our findings suggest that SNR of spoken sentences has no robust effect on the excitability of the articulatory motor cortex.

The present results highlight that the articulatory motor cortex is involved in speech processing both in optimal and challenging conditions. There is a growing body of evidence that the speech motor areas play a causal role in the processing of speech sounds. Indeed, several studies have shown that TMS-induced modulation of motor areas influence performance in demanding speech discrimination tasks, in which syllables were presented in noise or close to the category boundary to increase task difficulty (D’Ausilio et al., 2009; Meister, Wilson, Deblieck, Wu, & Iacoboni, 2007; Möttönen & Watkins, 2009; Smalle, Rogers, & Möttönen, 2015). Moreover, Schomers, Kirilina, Weigand, Bajbouj, & Pulvermüller (2015) showed that TMS over the tongue and lip motor representations in the left primary motor cortex affected reaction times in a word-to-picture matching task. Since the words were presented without noise in this study, these findings provide evidence that the articulatory motor system contributes to processing of meaningful speech in optimal listening conditions. Furthermore, TMS-induced disruptions in the articulatory motor cortex have been shown to modulate the processing of clear syllables in the auditory cortex (Möttönen, Dutton, & Watkins, 2013; Möttönen, Ven, & Watkins, 2014). Altogether, these findings are in line with the present results and demonstrate that the articulatory motor regions play a causal role in processing clear speech as well as degraded speech. However, it is possible that the contribution of motor areas to speech perception is greater in challenging conditions than in optimal listening conditions. This is not a trivial question to address experimentally, because degrading speech sounds increases task difficulty and consequently sensitivity to subtle motor manipulations (e.g., TMS-induced disruptions
in the motor system, see Möttönen & Watkins, 2012; Schomers & Pulvermüller, 2016). It is possible that some studies using clear speech stimuli have not been sensitive enough to measure the effect of motor manipulation (D’Ausilio, Bufalari, Salmas, & Fadiga, 2012; Sato, Tremblay, & Gracco, 2009).

In conclusion, the present results show that processing of ecologically valid speech signals (i.e., spoken sentences) in the articulatory motor system is robust across a wide range of SNRs and across coherent and anomalous semantic context. This demonstrates that the articulatory motor system is involved in speech perception both in optimal and in challenging listening conditions. The results do not support the idea that the recruitment of the articulatory motor cortex would increase in challenging listening conditions.

## Acknowledgements

We thank Dr. Matt Davis and Dr. Ingrid Johnsrude for the stimulus material. We are also grateful to the participants for contributing with their time and effort to this study. This study was funded by the Wellcome Trust (WT091070AIA) and Medical Research Council, UK (G1000566).

